# Complex+: Aided Decision-Making for the Study of Protein Complexes

**DOI:** 10.1101/744656

**Authors:** Mehrnoosh Oghbaie, Petr Šulc, David Fenyö, Michael Pennock, John LaCava

## Abstract

Proteins are the chief effectors of cell biology and their functions are typically carried out in the context of multi-protein assemblies; large collections of such interacting protein assemblies are often referred to as interactomes. Knowing the constituents of protein complexes is therefore important for investigating their molecular biology. Many experimental methods are capable of producing data of use for detecting and inferring the existence of physiological protein complexes. Each method has associated pros and cons, affecting the potential quality and utility of the data. Numerous informatic resources exist for the curation, integration, retrieval, and processing of protein interactions data. While each resource may possess different merits, none are definitive and few are wieldy, potentially limiting their effective use by non-experts. In addition, contemporary analyses suggest that we may still be decades away from a comprehensive map of a human protein interactome. Taken together, we are currently unable to maximally impact and improve biomedicine from a protein interactome perspective – motivating the development of experimental and computational techniques that help investigators to address these limitations. Here, we present a resource intended to assist investigators in (i) navigating the cumulative knowledge concerning protein complexes and (ii) forming hypotheses concerning protein interactions that may yet lack conclusive evidence, thus (iii) directing future experiments to address knowledge gaps. To achieve this, we integrated multiple data-types/different properties of protein interactions from multiple sources and after applying various methods of regularization, compared the protein interaction networks computed to those available in the EMBL-EBI Complex Portal, a manually curated, gold-standard catalog of macromolecular complexes. As a result, our resource provides investigators with reliable curation of bona fide and candidate physical interactors of their protein or complex of interest, prompting due scrutiny and further validation when needed. We believe this information will empower a wider range of experimentalists to conduct focused protein interaction studies and to better select research strategies that explicitly target missing information.

## Introduction

Protein complexes are assemblies of proteins within multi-component macromolecules. They are commonly defined by stable interactions that are likely to be observed in homeostatic cells; protein complexes are nevertheless understood to frequently contain labile and/or dynamical components that may not be apparent in the steady state. These stable and dynamical macromolecules are the prime effectors of cell biology. Thus, identifying the compositions and topologies of protein complexes from raw and partially processed protein-protein interactions (PPI) data is an important area of research. Because mutations in protein coding genes can alter the protein product, including changes to protein length, amino acid sequence, expression level, and subcellular localization, such changes can lead to the formation of altered PPI networks (a.k.a. interactomes) that cause cellular dysfunction and disease [1–4]. Hence, to understand health and disease, we must understand their associated interactomes. However, comprehensive interactome mapping has proven challenging for at least three reasons: 1) experiments to directly identify protein interactions are relatively expensive and 2) time consuming, e.g. compared to indirect genomic/transcriptomic methods; moreover 3) any single proteomic approach is likely to miss many PPIs (high false negatives) and incorrectly assign many spurious PPIs (false positives) [5–8].

Currently there are several well-known biological single/multiple protein search engines derived from interaction data as well as protein complex search engines that are manually curated from literature. Here we summarize a few of the most well-developed resources to contextualize our own offering.

- **STRING** [10] is arguably the most famous protein interaction search engine. STRING integrates different properties of molecular interactions including experimental repositories, computational prediction methods, and public text collections, to calculate a final link score and map them to a single interaction network: It is a single protein search engine.
- **GeneMania** [11] is single/multi-protein search engine: GeneMania [12] is known to be more refined compared to other protein search engines. It gathers data from a very limited number of publications and maps them to a single gene network to predict gene functions. Among protein search engines listed here, only GeneMania accepts multiple gene entry and extends the list with functionally similar genes.
- **BioPlex** [13, 14] is a single protein search engine with physical interactions, currently cataloging 5900 immunoprecipitation-mass spectrometry experiments; this was achieved using one cell type (HEK-293T), ectopically expressing a large number of affinity tagged human open reading frames.

There are also two search engines for finding protein complexes:

- **Complex Portal** [15, 16] is a manually selected resource of macromolecular complexes. All complexes are derived from physical molecular interaction evidence extracted from the literature and cross-referenced in IntAct [17], or by curator inference from information on homologs in closely related species. The search engine requires a complex name or complex ID and returns a network of protein complex interactions, structure, and functional information.
- Similar to Complex Portal, Corum [18] is a manually curated database of experimental data from mammalian protein complexes.

Despite these resources there remains a disconnect between tools that populate interaction networks and sources of manually curated knowledge about protein complexes: as of now, the protein complex search engines do not identify and retrieve putative and candidate interactors from the remaining, as-yet-un-curated data. Part of the difficulty of identifying potential members of a complex is that the search space of possible interactions is understood to be extremely large [2, 9], albeit not definitively quantified as yet. In light of this, and the above-stated challenges, the objective of this research is to aid molecular biologists by leveraging existing collections of experimental data and applying machine learning techniques to identify probable but yet unconfirmed interactions (candidate complex membership). Doing so will enable researchers to better target their experimental hypotheses and consequently, reduce the number of experimental trials required. We achieve this by applying network-based, random walk statistical algorithms to integrate existing heterogeneous data sets that describe various properties of PPIs. Unfortunately, manually curated data sets that describe bona fide physiological interaction networks are few, and such databases grow very slowly.

However, the algorithm can be trained using this curated data to make appropriate inferences from more voluminous but lower quality automatically curated data sets. For this research, data from the “gold standard” database, Complex Portal, were used to train the algorithm. Once trained, the algorithm can be used to predict which unconfirmed interactions from the lower quality data are probable candidates for verification via targeted experiments; we computationally validated this application by using Leave-One-Out Cross Validation (LOOCV) to assess the accuracy of the algorithm in predicting the current members of a complex. Therefore, we demonstrate that by appropriately merging the two types of data resources, the false-positive prediction rate can be reduced while promising new candidates can be effectively distinguished from noise focusing downstream experimental strategies and improving biological understanding.

### Use Case

In our interface, a typical use case includes feeding a network with the current understanding of a protein complex, including its known members and ontological associations (e.g. as defined by EMBL-EBI Complex Portal), in order to retrieve additional candidate members through expanding the network as follows:

1. Choosing the species of interest (only human has been extensively tested by us up to now).
2. Selecting the target protein complex using the ‘protein complex tab’ in the main panel -consequently, members of the selected complex will appear in the sidebar.
3. Choosing the number of additional candidate interactors to be shown in the aiding interface.
4. Optionally, adding other proteins to the target complex membership, in the case of un-curated evidence or private knowledge of their membership.
5. Calculating discriminant scores^1^ using the information fed to the algorithm. The algorithm will automatically reduce the size of the network to those proteins that are part of the cell component ontological annotation of the target protein complex.
6. Checking the result in the ‘interactions network tab’ in the main panel as shown in Figure 1.
7. Reviewing the studies that the support the inclusion of the additional candidate complex members retrieved.

**Fig 1.**
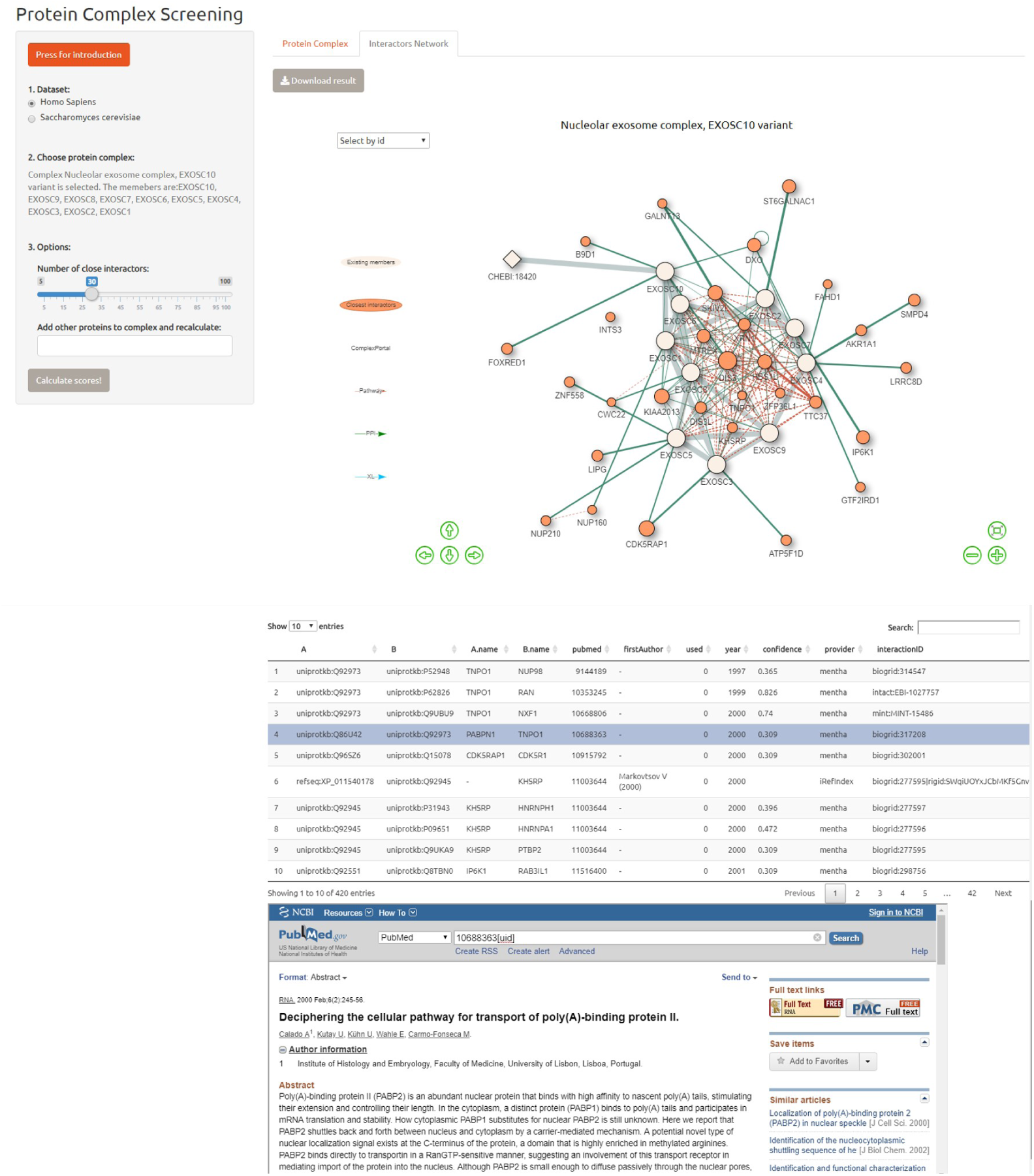
Interface design: View of the closest interactors networks.

## Materials and Methods

The approach to infer potential members of a protein complex is to fuse data from multiple heterogonous data sets including coexpression, colocalization, and cross-linking among others.To translate all the information to the protein level, we mapped networks with heterogeneous nodes into protein nodes using BiomaRT [38] and PaxDB [39]. Then, we applied ridge regression to weight all the sub-networks using existing protein interactions in Complex Portal as dependent variable. Once the weighted networks are complete, protein-protein interactions are predicted using networking scanning algorithms. Each of these components is described in more detail below.

### Data Collection

Different properties of human protein interactions were gathered as shown in Figures 1 & 2:

1. **Coexpression** -Two genes are connected if their expression levels correlate across conditions in a gene expression study. We used data from GeneMania which collected data associated with a publication from the Gene Expression Omnibus (GEO) [25].
2. **Colocalization** -Genes expressed in the same tissue, or proteins found in the same location. Two genes are linked if they are both expressed in the same tissue or if their gene products are both identified in the same intracellular location. We extract colocalization data from GeneMania.
3. **Prediction** -Predicted functional relationships between genes by mapping known functional relationships from another organism via orthology. We used GeneMania prediction data.
4. **Physical Interaction** -Two gene products are linked if they were found to interact in primary research; protein-protein interactions are store in databases such as BioGRID [26] and IntAct. Some of these records use gene name or uniprot ID to represent nodes. As the result, we had two layers of physical interaction data (one with genes as nodes and another one with uniprot IDs).
5. **Pathway Interaction** -represents protein-protein interactions observed within a functional pathway, extracted from databases such as: Reactome [27], NCI [28] and Panther [30].
6. **Cross-Linking** -Chemical protein cross-linking with analysis by mass-spectrometry is a method used to extract structural information about protein interactions and protein complexes at the peptide-peptide contact level. The output describes the cross-linked residues in two peptides and their proximity; We gathered our data from XlinkDB [31]. Cross-linking network links are binary in our network, and each edge represents a link with distance less than 25 Angstrom.
7. **Disease Similarity** -We used a previously described method [34] to calculate the disease similarity based on common phenotypes using disease records from OMIM [32] as well as phenotype annotation and graph structure from Human Phenotype Ontology (HPO) [27]. We only considered the five closest diseases in our network using the k-nearest neighbors algorithm.
8. **Domain Interaction** -Classifying protein sequences by their functional analysis into families and predicting the presence of domains and important sites [37]. Domain interactions are considered useful in predicting the ability of two proteins to interact within the context of a pathway or gene ontology term.

**Fig 2.**
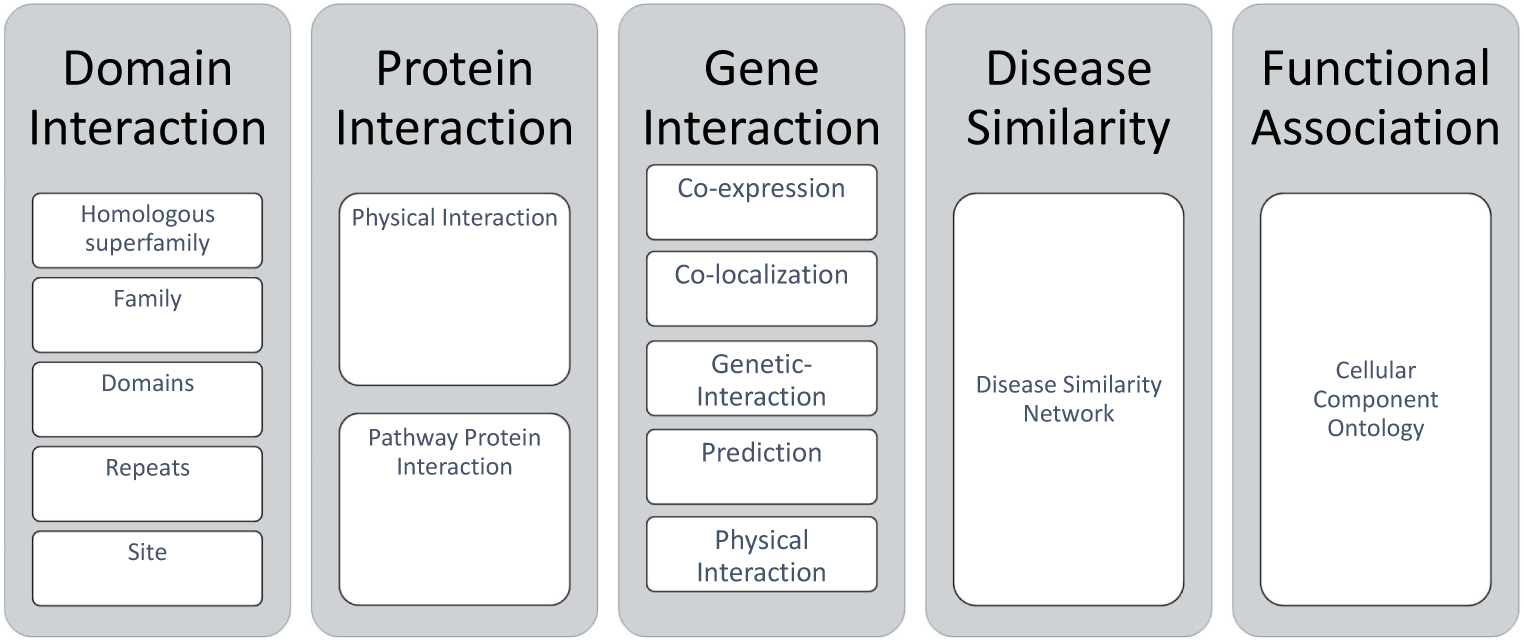
Data sources. Data gathered from different levels of abstraction.

### Mapping

In cases where multiple links exist between two nodes, we adopt the maximum score. When integrating heterogeneous data we initially used the assumption that different types of nodes map to one another in a binary fashion. For example, the assumption that a gene could produce all of its protein products with equal probability. This rational was applied to map connections from gene-level and disease-similarity networks to protein networks. BiomaRt [38] was initially used to construct this bipartite mapping between these networks. However, our analysis showed that such an assumption would increase the error and false positive rate. To overcome this problem, we used the PaxDB [39] database to calculate the protein abundance ratio of proteins resulting from the same gene. PaxDB is a whole proteome abundance database across organisms and tissues. Yet, we have other data layers that do not result from gene or protein interaction, and we could not find any method to optimally integrate them at this time.

### Network Scanning Algorithms

The random walk algorithm with restart parameter and the label propagation algorithm (LPA) have been well-described in the literature [19–21]. The basic idea of the random walk algorithm is that we start a metaphorical walker at a node that is known to be a member of a protein complex. This walker then traverses the network by randomly following different links in the network. If we repeat the process with many walkers, we can determine which other proteins can be reached from the starting protein. We can also estimate the probability that each of these proteins will be visited by a walker. It is natural to assume that local neighbors of the starting node would be visited more often. LPA [22] is a generalization of the local neighborhood approach where a discriminant score is calculated by the sum of weighted scores of its direct interactions.

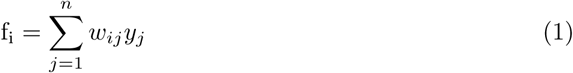

If you iterate discriminant scoring to n neighbors, it will converge to the random walk. In a restart random walk, a random walker will stay in the initial node with a probability of r and go to other nodes with probability of 1-r. The RRW algorithm has been used in proteomic studies and is known to perform best with r being set at 0.7-0.75 [20, 23, 24].

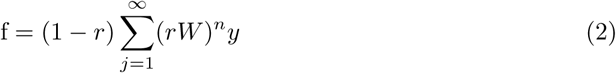

Under the condition that *rW ≤* 1 and 0 *< r <* 1, the infinite sum in Eq.2 could be simplified to Eq.5. We implemented symmetric normalization demonstrated in Eq. 4 to guarantee that *rW ≤* 1 is valid. Previous literature [29] extensively investigated the performance of LPA with different normalization methods and noted that symmetric normalization had the highest precision among all.

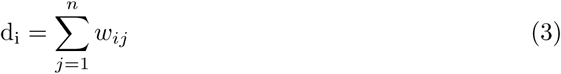

Where *D* is the diagonal matrix of row sums of the weighted matrix, the normalized symmetric version of the weighted matrix 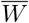 replaces *W* in the Eq.5.

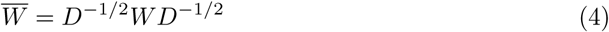

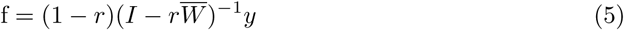

### Multiple Kernel Network

An identified link between two proteins represents an assembly of incomplete biological significance. It is often required to integrate data from different properties into a single network. Having d normalized networks, 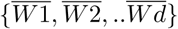, we want to construct a single weighted networks 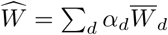 with *α*_*d*_ *>* 0 that is optimized to predict the existence of interactions. If we try to solve this problem using linear regression, our regression equation could be written as:

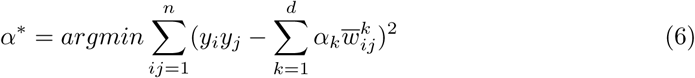

Where *y*_*i*_ is the output vector of node i and 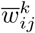 represents the edge between node *i* and *j*. To find a vector of network weights that minimizes the cost function, we used ridge regression. Ridge regression is a technique that is normally used when there is multicollinearity among different variables or there is correlation between independent variables. When multicollinearity occurs, least squares estimates are small but their variances are large values and may be far from the true values.Using a fast assumption from the previous literature [11] [29], we calculated *α*_*d*_ as follow:

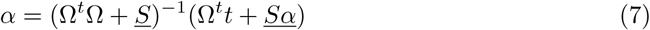

Where *<u>S</u>* is the precision matrix and calculated as diagonal matrix 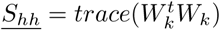, Ω is a matrix of *n*^2^x*k* that each column is the vectorized version of 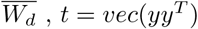 and *<u>α</u>* is set to constant value 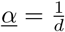.

## Result

Ridge regression was used to integrate multiple networks into a single network; stable links were predicted using a label propagation algorithm. We incrementally added the network layers and used Leave-One-Out Cross validation (LOOCV). LOOCV is a special case of Leave P Out Cross-validation with p=1 whereby, during each run, statistics are gathered on the sample that was ‘left-out’. In a typical scenario an input vector, e, represents protein complex members, where the indices corresponding to the k protein complex members are assigned a value of 1. In the first step, one of the k proteins will be ‘left out’ and the corresponding index in vector e will be changed to 0. Using the restart random walk formula and parameter, r, discriminant scores of all the nodes will be calculated as follow:

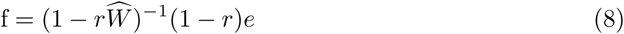

After leaving out the calculated discriminant scores of k-1 indices that had a value of 1 in the input vector, we calculated the rank of the left out protein. The process has to be repeated k times to gather statistics for all the protein members of a complex. The lower the rank of a protein, the better the performance of the algorithm in predicting the complex membership of that protein. The process can be segregated into the following steps:

i. We trained our network by (1) randomly removing 10% of our records (protein complexes in Complex Portal) (2) building a matrix containing the remaining 90% of the records and (3) measuring the weight of layers using Ridge Regression as explained in the previous section. Then we repeated the three steps 10 times. Finally, we calculated the average weight of each layer in all runs. Note, the results, displayed in Figure 4, indicate our approach is agnostic to 10% random data loss.
ii. We constructed different combination of layers^2^ to compare their performance as follows:
  - PPIs ^3^
  - Biological pathways
  - PPIs + protein cross-linking
  - PPIs + biological pathways
  - PPIs + protein cross-linking + biological pathways
  - PPIs + protein cross-linking + biological pathways + genetic interactions ^4^
  - PPIs + protein cross-linking + biological pathways + disease similarities
  - PPIs + protein cross-linking + biological pathways + disease similarities + genetic interactions
iii. LOOCV was carried out on all the protein complexes to compare the performance of different combinations of layers. Each time, one protein was left out from a target complex and the rest of the proteins were used as a seed to find the closest interactors in the network using the restart random walk algorithm.
iv. Finally, a cumulative distribution of the ranks was plotted to determine the contribution of each layer when predicting the known members of a protein complex, permitting the evaluation of the performance of different combinations of layers.

We observed the following results: (i) the network layers associated with biological pathways, followed by PPIs, contained the most useful information; (ii) the protein cross-linking network layer did not contribute much information -which is not unexpected considering the sparsity of data; (iii) the disease similarities network layer appeared to reduce the accuracy of algorithm; and finally, (iv) the optimum scenario observed during our testing can be attributed to a combination of PPIs, biological pathways, protein cross-linking, electronic prediction, colocalization, and coexpression network layers.

**Fig 3.**
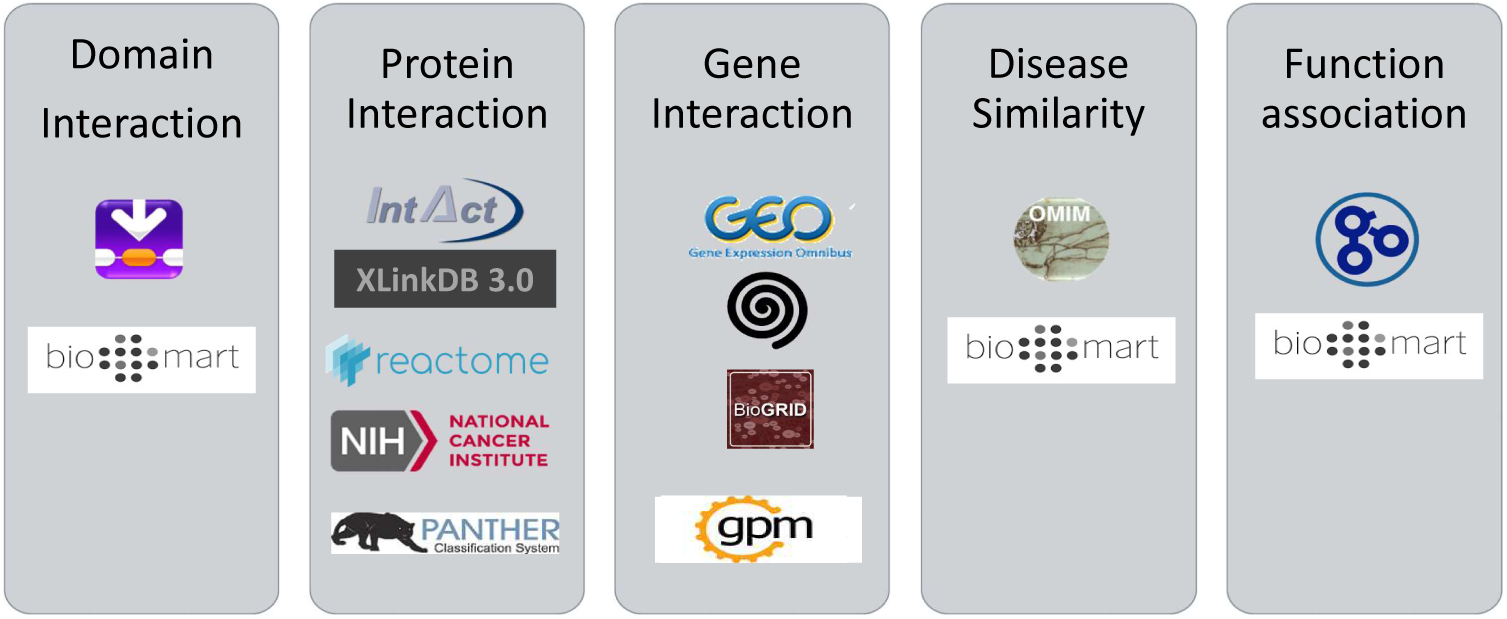
Data mapping. Data gathered from different sources: biomart and paxDB were used for mapping between layers.

**Fig 4.**
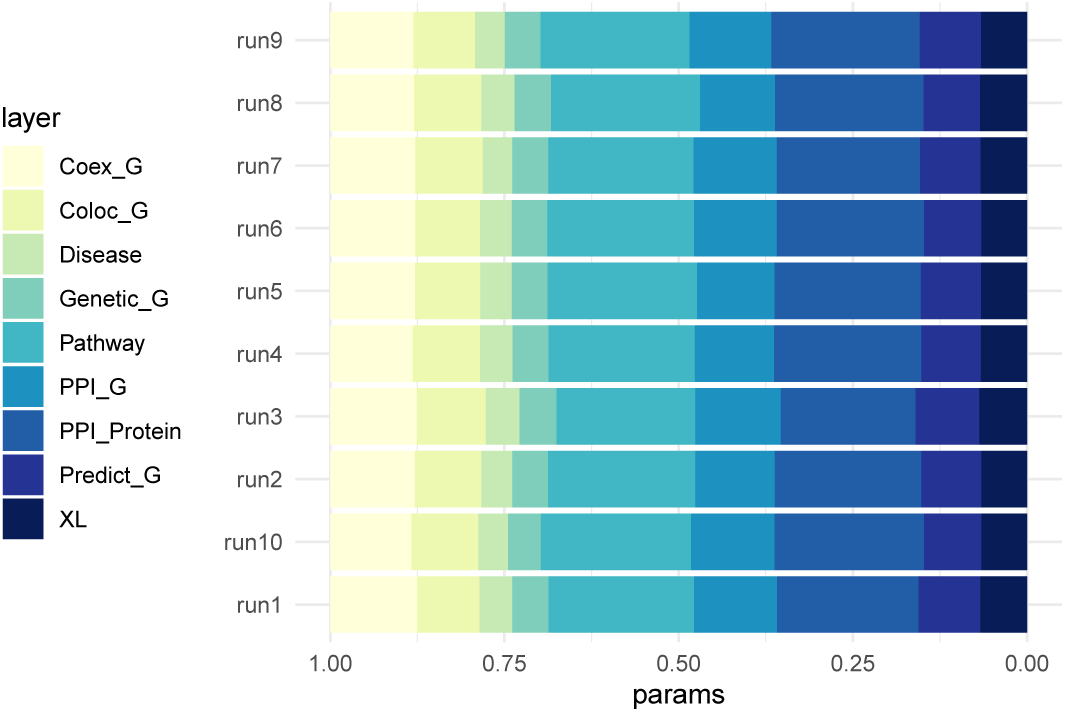
Validation result. Validation result from 10 trials.

**Fig 5.**
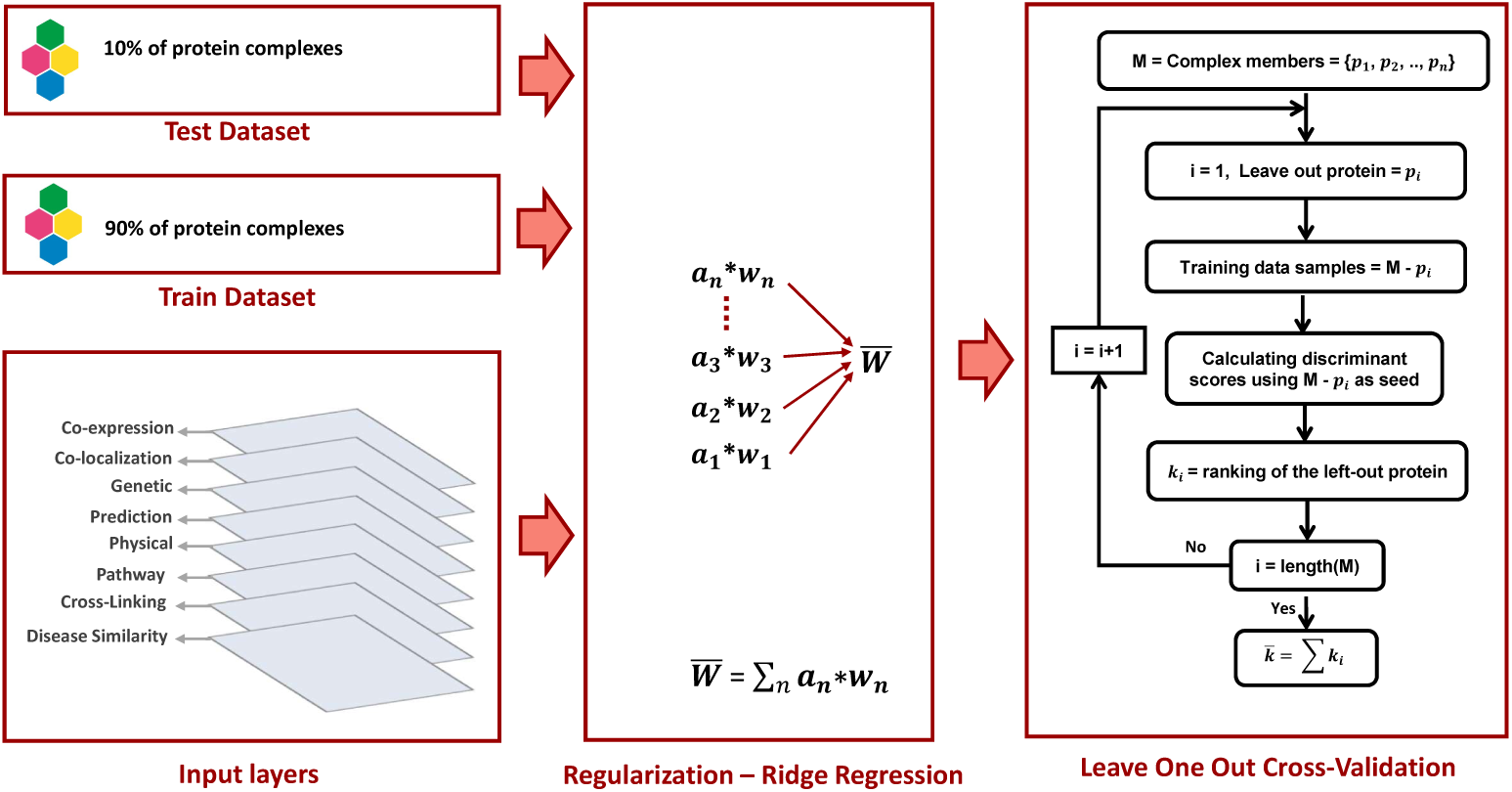
Validation pipeline.

**Fig 6.**
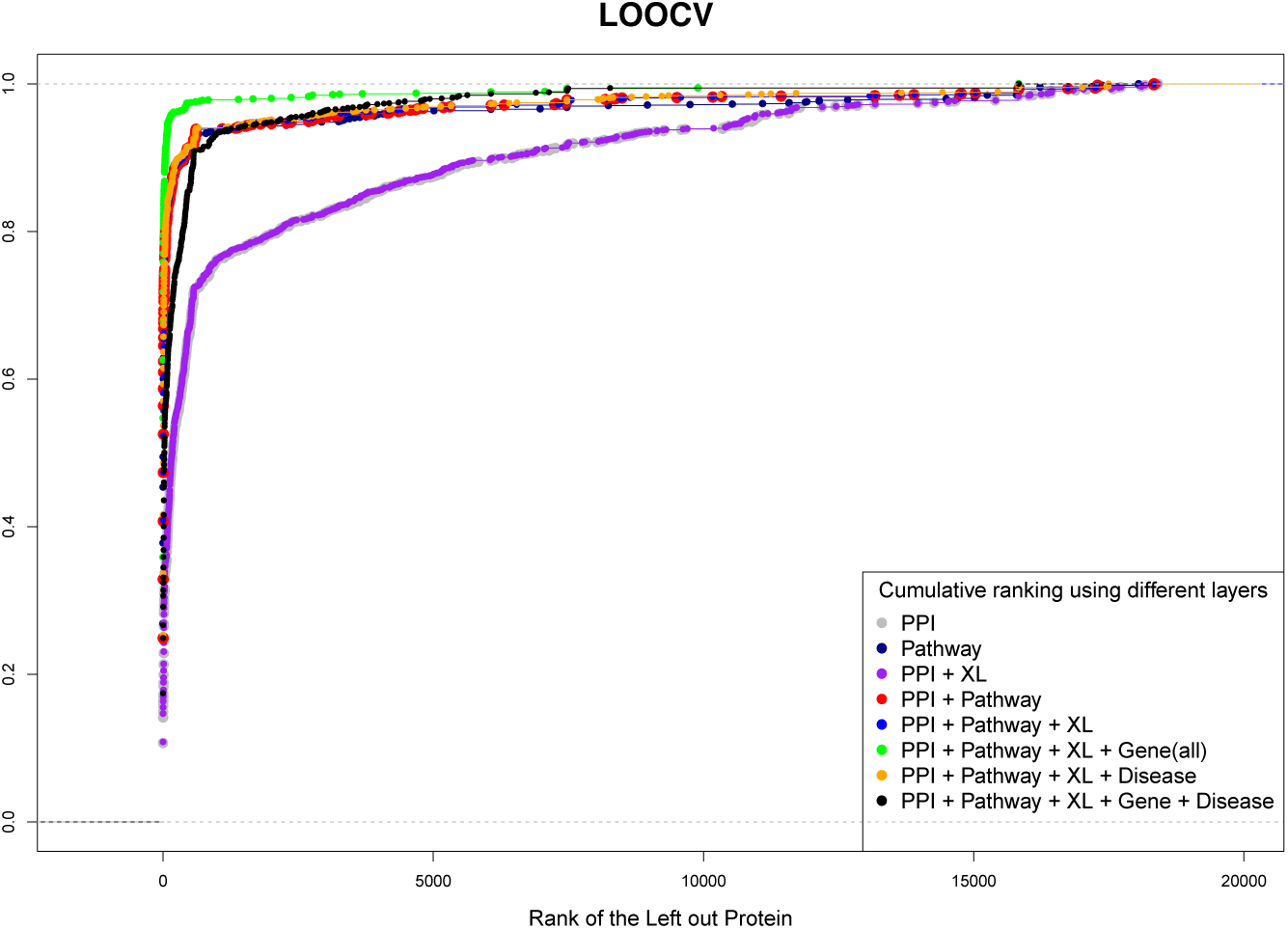
Comparing different combination of layers. Leave One Out Cross-validation result.

## Discussion

Mapping different layers of information is laborious due to the non-linear nature of connections between them and known false positive detection rate in interactomic databases. LOOCV showed that disease similarities network did not positively contribute to detecting the current members of protein complexes when aggregated with other layers into a monoplex protein interaction network. It is possible that implementing disease similarities with the rest of the layers as a multiplex network and filtering disease similarities to those the target complex is known to interact with or contribute to (Complex Portal represents diseases with their Orphanet ID) could improve the performance of the algorithm as was investigated in previously [34]. This would require feeding the algorithm with the list of protein members and the diseases they are associated with. Although protein cross-linking data did not prove to be of abundant value in the current study, we are hopeful that its value will increase with data size and that the peptide-peptide contacts provided will obviate the need for other forms of structural information (except for in cross-validation). Future work will focus on: (i) wet lab validations of a select putative candidate members of protein complexes produced by our algorithm; (ii) extending the complex membership prediction algorithm to also contain RNA-protein and DNA-protein interfaces aggregated from available resources (PDB and NDB databases [41]); and (iii) extending this method to numerous organisms.

The number of available protein complex records in a manually curated database such as Complex Portal is limited and so is the average number of links in curated complexes (less than 5). Consequently, it is implausible to tune each complex by matching it to other complexes with similar functional associations. In this paper, we presented a tool to predict potential members of a protein complex using heterogeneous data sets integrated into a single network. We tuned our network using the total available links as one entity. While we were not able to integrate all the available information (such as disease similarity, and structural information from domains), we still achieved robust predictive results when we applied LOOCV. In fact, the result does not seem to be sensitive to losing 10% of the links. We believe that it is necessary to study proitein complexes within the context of their broader interaction networks. We acknowledge the divide between an ideal protein complex search engine and present resources to describe protein complexes, including those presented here; but we believe our approach can be of significant utility to molecular biologists aiming to close potentially non-obvious knowledge gaps in interactome-related studies. Considering the dynamic nature of the interactome, providing more data sources and features in the interface is necessary and will be pursued in future versions.

## Supporting information

Project code is available at https://github.com/moghbaie/ComplexPlus

## Acknowledgments

This research was supported in part by NIH grant R01GM126170 to J.L.

## Footnotes

1 Discriminant score is calculated by aggregating weighted score of all direct interactions of a node

2 Different layers of data and their sources are discussed in Data Collection section

3 including both gene and protein physical interaction

4 coexpression, colocalization, and electronic prediction

## References

1. Sharma, A. et al. A disease module in the interactome explains disease heterogeneity, drug response and captures novel pathways and genes in asthma. Hum. Mol. Genet. 24, 3005–3020.

2. Menche, J. et al. Disease networks. Uncovering disease-disease relationships through the incomplete interactome. Science 347, 1257601 (2015).

3. Barabasi, A.-L., Gulbahce, N. Loscalzo J. Network medicine: a network-based approach to human disease. Nat Rev Genet 12, 56–68 (2011).

4. Harris, C. C. Protein-protein interactions for cancer therapy. Proc Natl Acad Sci USA 103, 1629–1660 (2006).

5. Hart GH, Ramani AK, Marcotte EM. How complete are current yeast and human protein-interaction networks?. Genome Biol. 2006. doi: 10.1186/gb-2006-7-11-120.

6. Hakhverdyan, Z. et al. Rapid, optimized interactomic screening. Nat Methods 12, 553–560 (2015).

7. Mellacheruvu, D. et al. The CRAPome: a contaminant repository for affinity purification-mass spectrometry data. Nat Methods (2013).

8. Boulon, S. et al. Establishment of a protein frequency library and its application in the reliable identification of specific protein interaction partners. Mol Cell Proteomics 9, 861–879 (2010).

9. Stumpf, M. P. H. et al. Estimating the size of the human interactome. Proc Natl Acad Sci USA 105, 6959–6964 (2008).

10. Franceschini A, Jensen LJ et al. STRING v9.1: protein-protein interaction networks, with increased coverage and integration. Nucleic Acids Research. 2013 Jan. doi: 10.1093/nar/gks1094.

11. Sara Mostafavi, Debajyoti Ray, David Warde-Farley, Chris Grouios and Quaid Morris. GeneMANIA: a real-time multiple association network integration algorithm for predicting gene function. Genome Biology. 2008 June;https://doi.org/10.1186/gb-2008-9-s1-s4.

12. Rintaro Saito, Trey Ideker et al. A travel guide to Cytoscape plugins. Nature Methods 9, pages 1069–1076 (2012).

13. Huttlin EL, Gygi SP, Harper JW et al. Architecture of the human interactome defines protein communities and disease networks. Nature. 2017 May. doi: 10.1038/nature22366.

14. Huttlin EL, Gygi SP et al. The BioPlex Network: A Systematic Exploration of the Human Interactome. Cell. 2015 Jul 16. doi: 10.1016/j.cell.2015.06.043.

15. Meldal BH, Orchard S et al. The complex portal–an encyclopaedia of macromolecular complexes. Nucleic Acids Res. 2015 Jan 2015. doi: 10.1093/nar/gku975.

16. Birgit HM Meldal and et al. Complex Portal 2018: extended content and enhanced visualization tools for macromolecular complexes. Nucleic Acids Research. 2018 Oct 24. doi: 10.1093/nar/gky1001.

17. Samuel Kerrien, Henning Hermjakob and et al. The Intact molecular interaction databse in 2012. Nucleic Acids Research. Volume 40, Issue D1, 1 January 2012, pages D841–D846.

18. Madalina Giurgiu, Andreas Ruepp et al. CORUM: the comprehensive resource of mammalian protein complexes—2019. Nucleic Acids Research, 2019 January 08. doi: 10.1093/nar/gky973.

19. Sebastian Köhler, Sebastian Bauer, Denise Horn, Peter N. Robinson. Walking the Interactome for Prioritization of Candidate Disease Genes. AJHG, 2008 Apr 11. doi: 10.1016/j.ajhg.2008.02.013

20. Kathy Macropol, Tolga Can, Ambuj K Singh. RRW: repeated random walks on genome-scale protein networks for local cluster discovery. BMC Bioinformatics, 2009 Sep 09. doi: 10.1186/1471-2105-10-283

21. Osamu Maruyama, Ayaka Chihara. NWE: Node-weighted expansion for protein complex prediction using random walk distances. Proteome Science, 2011 Oct 14. doi: 10.1186/1477-5956-9-S1-S14

22. Marcotte EM, Pellegrini M, Thompson MJ, Yeates TO, Eisenberg D.. A combined algorithm for genome-wide prediction of protein function. Nature, 1999 Nov 4. doi: 10.1038/47048

23. Chen X, Liu MX, Yan GY. Drug-target interaction prediction by random walk on the heterogeneous network. Mol Biosyst. 2012 Jul 6. doi: 10.1039/c2mb00002d.

24. Navlakha S, Kingsford C. The power of protein interaction networks for associating genes with diseases. Bioinformatics. 2010 Apr 15. doi: 10.1093/bioinformatics/btq076.

25. Barrett T, Soboleva A et al. NCBI GEO: archive for functional genomics data sets–update. Nucleic Acids Res. 2013 Jan. doi: 10.1093/nar/gks1193.

26. Oughtred R, Tyers M et al. The BioGRID interaction database: 2019 update. Nucleic Acids Res. 2019 Jan. doi: 10.1093/nar/gky1079.

27. Antonio Fabregat, Henning Hermjakob et al. Reactome pathway analysis: a high-performance in-memory approach. BMC Bioinformatics. 2017. doi:10.1186/s12859-017-1559-2.

28. Carl F. Schaefer, Kira Anthony, Shiva Krupa, Jeffrey Buchoff, Matthew Day, Timo Hannay, Kenneth H. Buetow. PID: the Pathway Interaction Database. Nucleic Acids Res. 2009 Jan. doi:10.1093/nar/gkn653.

29. Sara Mostafavi. Computational Prediction of Gene Function from High-throughput Data Sources. thesis for the degree of Doctor of Philosophy Graduate Department of Computer Science University of Toronto, 2011.

30. Huaiyu Mi, Anushya Muruganujan, Paul D. Thomas. PANTHER in 2013: modeling the evolution of gene function, and other gene attributes, in the context of phylogenetic trees. Nucleic Acids Res. 2013 Jan. doi:10.1093/nar/gks1118.

31. Juan Chavez, CF Lee, A Caudal, Andrew Keller, R Tian, James Bruce. Chemical crosslinking mass spectrometry analysis of protein conformations and supercomplexes in heart tissue.. Cell Syst. 2018 Jan 24. doi: 10.1016/j.cels.2017.10.017.

32. Huaiyu Mi, Anushya Muruganujan, and Paul D. Thomas. Phenotypic and Genotypic Analyses of Genetic Skin Disease through the Online Mendelian Inheritance in Man (OMIM) Database. Nucleic Acids Res. 2013 Jan. doi: 10.1093/nar/gks1118.

33. Sebastian Köhler et. al The Human Phenotype Ontology project: linking molecular biology and disease through phenotype data. Nucleic Acids Res. 1 January 2014. doi: 10.1093/nar/gkt1026.

34. Valdeolivas A, Tichit L, Navarro C, Perrin S, Odelin G, Levy N, Cau P, Remy E, Baudot A. Random walk with restart on multiplex and heterogeneous biological networks. Bioinformatics. 2019 Feb 1 doi: 10.1093/bioinformatics/bty637.

35. Schweppe DK, Zheng C, Chavez JD, Navare AT, Wu X, Eng JK, Bruce JE. XLinkDB 2.0: integrated, large-scale structural analysis of protein crosslinking data. Bioinformatics. 2016 Sep 1. doi: 10.1093/bioinformatics/btw232.

36. Marc Vidal, Michael E. Cusick, Albert-Laszlo Barabasi. Interactome Networks and Human Disease. Cell. 2011 Mar 11. doi.org/10.1016/j.cell.2011.02.016.

37. Sarah Hunter, Corin Yeats et al. InterPro: the integrative protein signature database. Nucleic Acids Research, Volume 37, Issue Suppl 1, 1 January 2009, Pages D211–D215.

38. Steffen Durinck, Paul T Spellman, Ewan Birney, Wolfgang Huber. Mapping identifiers for the integration of genomic datasets with the R/Bioconductor package biomaRt. Nature. 2009 July 23.

39. Wang M, Herrmann CJ, Simonovic M, Szklarczyk D, von Mering C. Version 4.0 of PaxDb: Protein abundance data, integrated across model organisms, tissues, and cell-lines. Proteomics. 2015 Sep. doi: 10.1002/pmic.201400441.

40. Malay Kumar Basu, Eugenia Poliakov, Igor B. Rogozin. Domain mobility in proteins: functional and evolutionary implications. Briefings in Bioinformatics, Volume 10, Issue 3, May 2009, Pages 205–216, https://doi.org/10.1093/bib/bbn057.

41. Benjamin A. Lewis, Drena Dobbs et al. PRIDB: a protein–RNA interface database. Nucleic Acids Research, Volume 39, Issue Suppl 1, 1 January 2011, Pages D277–D282, https://doi.org/10.1093/nar/gkq1108.

